# Mild activation of the mitochondrial unfolded protein response increases lifespan without increasing resistance to stress

**DOI:** 10.1101/2024.09.13.612912

**Authors:** Alexa Di Pede, Bokang Ko, Abdelrahman AlOkda, Aura A. Tamez González, Jeremy M. Van Raamsdonk

## Abstract

The mitochondrial unfolded protein response (mitoUPR) is a stress response pathway that responds to mitochondrial insults by altering gene expression to recover mitochondrial homeostasis. The mitoUPR is mediated by the stress-activated transcription factor ATFS-1. Constitutive activation of ATFS-1 increases resistance to exogenous stressors but paradoxically decreases lifespan. In this work, we determined the optimal levels of expression of activated ATFS-1 with respect to lifespan and resistance to stress by treating constitutively-active *atfs-1(et17)* worms with different concentrations of RNA interference (RNAi) bacteria targeting *atfs-1.* We observed the maximum lifespan of *atfs-1(et17)* worms at full-strength *atfs-1* RNAi, which was significantly longer than wild-type lifespan. Under the conditions of maximum lifespan, *atfs-1(et17)* worms did not show enhanced resistance to stress, suggesting a trade-off between stress resistance and longevity. The maximum resistance to stress in *atfs-1(et17)* worms occurred on empty vector (0% *atfs-1* knockdown). Under these conditions, *atfs-1(et17)* worms are short-lived. This indicates that constitutive activation of ATFS-1 can increase lifespan or enhance resistance to stress but not both, at the same time. Finally, we determined the timing requirements for ATFS-1 to affect lifespan. We found that knocking down *atfs-1* expression only during development is sufficient to extend *atfs-1(et17)* lifespan, while adult-only knockdown has no effect. Overall, these results demonstrate that constitutively active ATFS-1 can extend lifespan when expressed at low levels and that this lifespan extension is not dependent on the ability of ATFS-1 to enhance resistance to stress.

## Introduction

The mitochondrial unfolded protein response (mitoUPR) is a stress response pathway that allows organisms to recover from various insults to the mitochondria ^1–3^. Transcriptional changes resulting from activation of the mitoUPR are mediated by the transcription factor ATFS-1/ATF5 (activating transcription factor associated with stress 1) ^4^ in combination with the transcription factor DVE-1 and the transcriptional regulator UBL-5 ^2^. ATFS-1 is normally localized to the cytoplasm where its mitochondrial targeting sequence (MTS) targets ATFS-1 to the mitochondria. At mitochondria, ATFS-1 is imported through the HAF-1 channel and degraded by the protease CLPP-1/CLPP ^4^. Under conditions of mitochondrial stress, the import of ATFS-1 into the mitochondria is inhibited. As a result, the nuclear localization signal of ATFS-1 targets cytoplasmic ATFS-1 to the nucleus to change gene expression in order to restore mitochondrial homeostasis. Among the target genes of the mitoUPR are mitochondrial chaperones, such as HSP-6 and HSP-60, which act to restore protein folding in the mitochondria ^5^.

The mitoUPR transcription factor ATFS-1 has been shown to affect both lifespan and resistance to stress. While deletion or knockdown of *atfs-1* does not decrease lifespan in wild-type worms ^6–8^, disruption of *atfs-1* can reduce the longevity of multiple long-lived mutant strains including *nuo-6* and *glp-1* worms ^8–10^, though this has not been observed under all conditions ^7^. This suggests that activation of ATFS-1 can contribute to lifespan extension in long-lived mutants.

However, constitutive activation of ATFS-1 in a wild-type background results in decreased lifespan ^7,11^. Interestingly, genes and interventions that activate the mitoUPR have been shown to increase lifespan, decrease lifespan or have no effect on longevity ^7,12,13^. This suggests the possibility that mitoUPR activation can contribute to longevity only under specific conditions.

Deletion of *atfs-1* decreases resistance to multiple exogenous stressors including oxidative stress, osmotic stress, heat stress and anoxia, and is required for the enhanced stress resistance of long-lived mitochondrial mutants ^8,11,14^. Consistent with a role for the mitoUPR in protecting against multiple stressors, we have found that constitutive activation of ATFS-1 results in increased resistance to oxidative stress, osmotic stress and anoxia ^11^, while others have reported that activation of ATFS-1 can protect against bacterial pathogen stress ^15^ and anoxia-reperfusion-induced death ^16^.

Given the strong correlation between stress resistance and longevity ^17,18^, it is somewhat surprising that constitutive activation of ATFS-1 results in decreased lifespan despite enhancing resistance to exogenous stressors. In addition, it has generally been observed that increased expression of the transcription factors that mediate stress response pathways can extend longevity, including DAF-16 ^19^, HSF-1 ^20,21^, SKN-1 ^22^, and HIF-1 ^23^. Interestingly, constitutive activation of XBP-1 only increases lifespan when expressed in specific tissues, while ubiquitous expression of constitutively active of XBP-1 decreases longevity ^24^. This result suggests the possibility that activation of stress response pathways might only increase lifespan under specific conditions.

In this work, we sought to determine whether activation of ATFS-1 could increase lifespan. To do this, we characterized the lifespan and stress resistance of constitutively active *atfs-1(et17)* mutants ^25^ when treated with different concentrations of *atfs-1* RNAi to decrease the levels of activated ATFS-1. We also examined the effect of decreasing the levels of constitutively active *atfs-1* only during development of only during adulthood on the lifespan of *atfs-1(et17)* mutants. We found that expression of a low level of constitutively active ATFS-1 extends longevity without enhancing resistance to stress.

## Results

### Decreasing expression of constitutively active ATFS-1 increases lifespan but reduces stress resistance

To determine whether constitutive activation of ATFS-1 could increase lifespan at lower levels and to identify the optimal dose of constitutively active ATFS-1 with respect to lifespan, we first examined the lifespan of worms heterozygous for the constitutive activation of *atfs-1.* This was accomplished by crossing *hsp-6p::GFP* worms to *atfs-1(et17)* worms and picking green fluorescent cross progeny to obtain *hsp-6p::GFP/+;atfs-1(et17)/+* animals. We then compared the lifespan of these animals to *atfs-1(et17)* homozygotes. We found that the lifespan of *atfs-1(et17)/+* animals is not statistically different from *atfs-1(et17)* homozygotes (**Figure S1**).

As an alternative approach, we treated the constitutively active *atfs-1(et17)* mutant with a dilution series of RNAi targeting *atfs-1* and measured lifespan. The dilution series included (1) 0% *atfs-1* RNAi + 100% empty vector (EV); (2) 25% *atfs-1* RNAi + 75% EV; (3) 50% *atfs-1* RNAi + 50% EV; (4) 75% *atfs-1* RNAi + 25% EV; and (5) 100% *atfs-1* RNAi + 0% EV (**Figure 1a**). For comparison, we also examined the effect of the *atfs-1* RNAi dilution series in wild-type worms.

**Figure 1.**
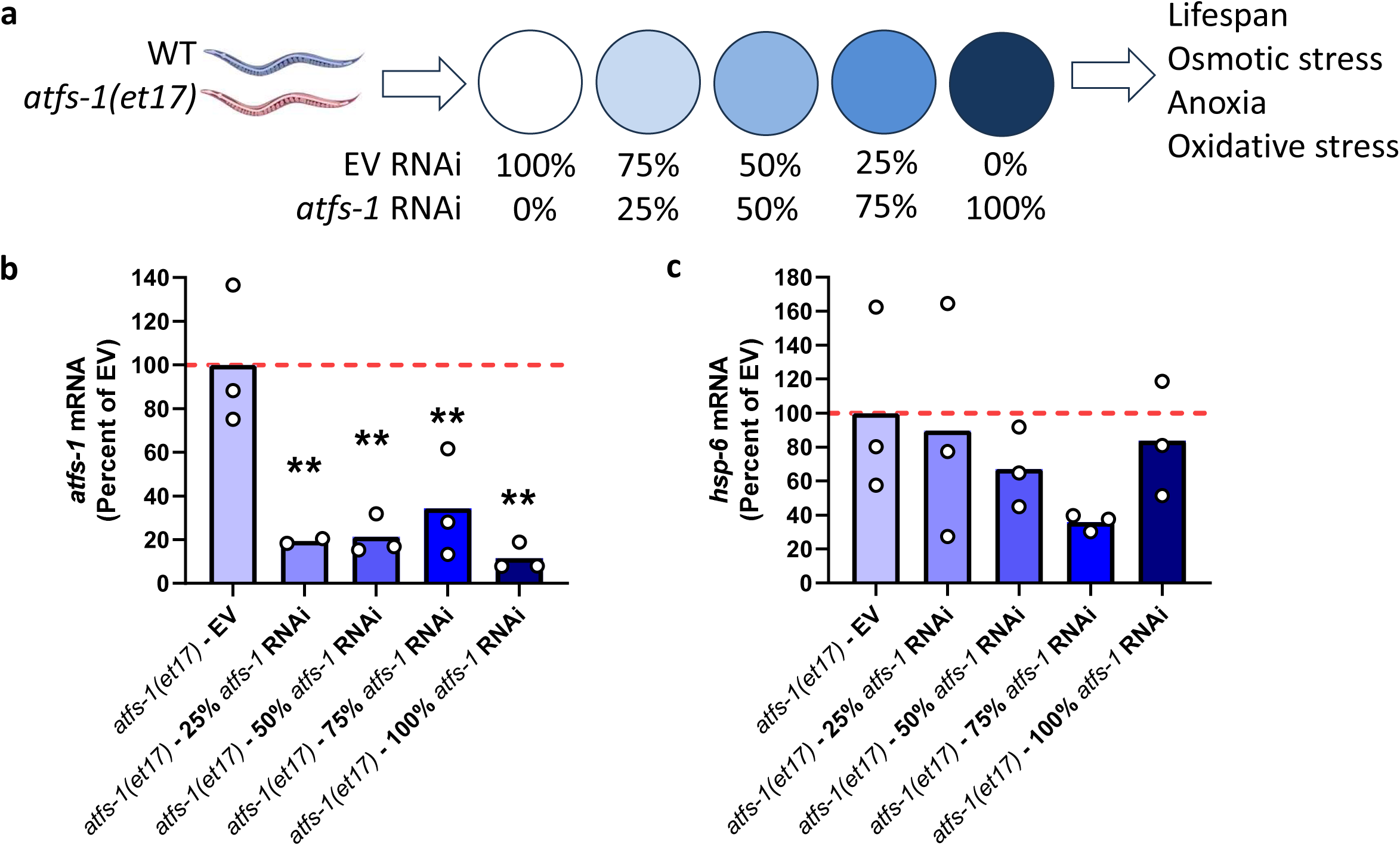
RNA interference targeting *atfs-1* decreases *atfs-1* and *hsp-6* mRNA levels in a dose-dependent manner. **a)** Wild-type and constitutively active *atfs-1(et17)* mutants were treated with different doses of *atfs-1* RNAi ranging from 0% to 100% (full strength). The effect of *atfs-1* RNAi on mRNA levels was measured using quantitative RT-PCR. **b)** *atfs-1* RNAi resulted in a significant decrease in *atfs-1* mRNA levels compared to worms treated with empty vector. **c)** Knockdown of *atfs-1* resulted in a trend towards decreased levels of *hsp-6* mRNA, which failed to reach significance. Three biological replicates were performed. Statistical significance was determined using a one-way ANOVA with Dunnett’s multiple comparisons test.

Before measuring lifespan, we used quantitative RT-PCR to confirm that treatment with *atfs-1* could effectively decrease *atfs-1* mRNA levels in *atfs-1(et17)* worms. We also measured the resulting effect on the ATFS-1 target gene *hsp-6.* RNAi treatment was initiated at the L4 stage of the parental generation and expression was measured when progeny reached young adulthood. We found that all four dilutions of *atfs-1* RNAi successfully decreased the levels of *atfs-1* mRNA (**Figure 1b**). Although *atfs-1* RNAi treatment resulted in a trend towards decreased *hsp-6* mRNA levels, these differences did not reach significance (**Figure 1c**). This could be due to variability or that the time point chosen was too early to observe the effects of *atfs-1* knockdown. It is also possible that the low level of *atfs-1* remaining after *atfs-1* knockdown is sufficient for normal *hsp-6* expression.

Having confirmed that the *atfs-1* RNAi effectively reduced the levels of *atfs-1*, we next examined the effect on lifespan. We found that *atfs-1* RNAi increased the lifespan of *atfs-1(et17)* worms (**Figure 2a**). While *atfs-1(et17)* worms lived shorter than wild-type worms on empty vector (EV; 0% *atfs-1* RNAi), these worms lived significantly longer than wild-type worms on full-strength 100% *atfs-1* RNAi, which was the concentration where we observed the maximum lifespan. Compared to *atfs-1(et17)* worms treated with empty vector (EV), even *atfs-1(et17)* worms treated with 25% *atfs-1* RNAi exhibited a significant increase in lifespan. In contrast, *atfs-1* RNAi did not affect the lifespan of wild-type worms at any concentration (**Figure 2b**).

**Figure 2.**
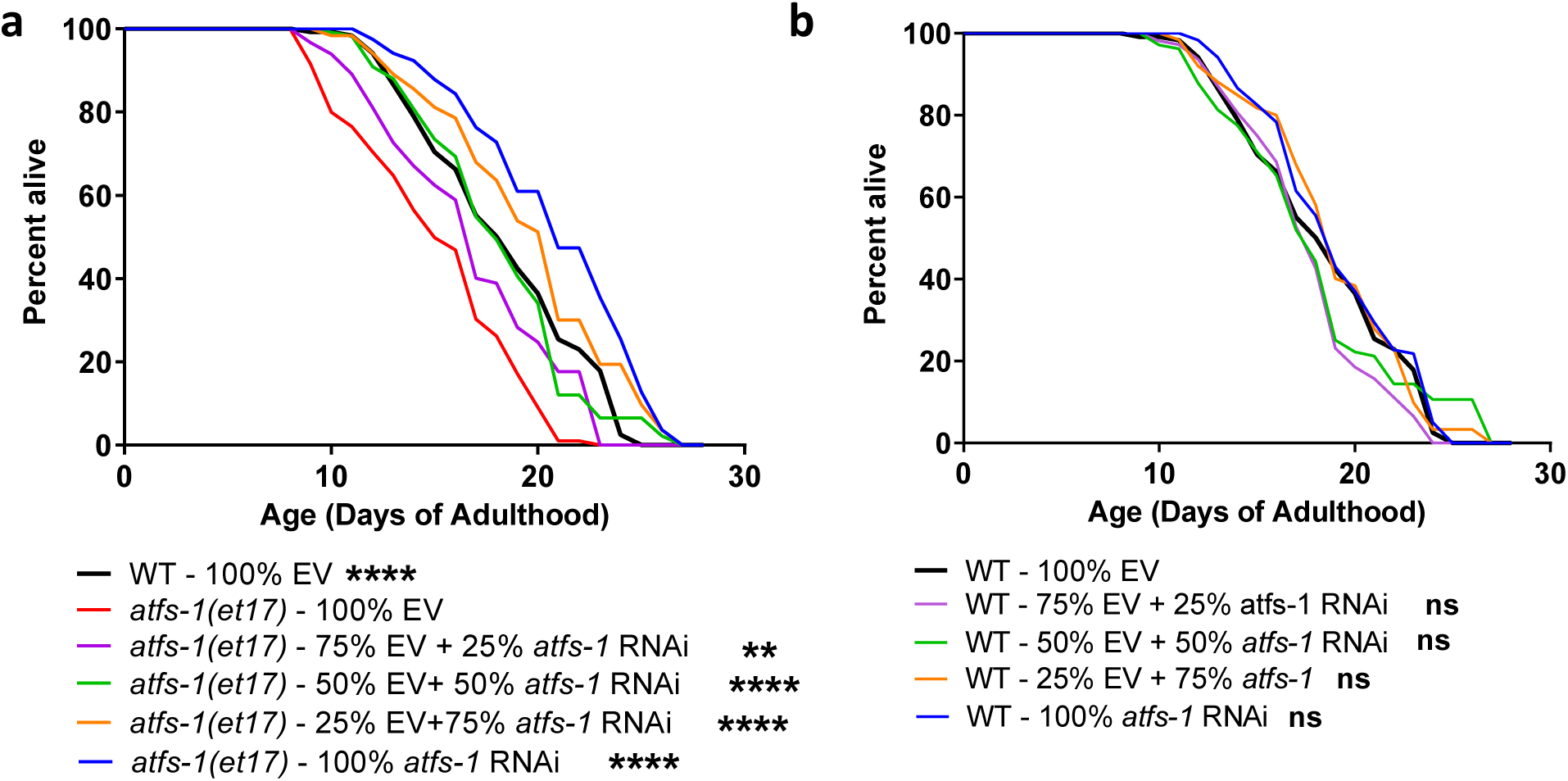
Low levels of constitutively active ATFS-1 increase lifespan. To determine the optimal level of constitutively active ATFS-1 with respect to lifespan, short-lived *atfs-1(et17)* mutants were treated with different concentrations of *atfs-1* RNAi ranging from 0% to 100%. (**a**) Compared to *atfs-1(et17)* worms on empty vector (EV), decreasing the expression of *atfs-1* with *atfs-1* RNAi increased lifespan with the maximum lifespan being observed at 100% *atfs-1* RNAi. *atfs-1(et17)* worms treated with 100% *atfs-1* RNAi live significantly longer than wild-type worms. (**b**) In contrast, *atfs-1* RNAi did not affect the lifespan of wild-type worms at any concentration. Three biological replicates were completed. Statistical significance was assessed using a log-rank test. Statistically significant differences from 100% EV group are shown. ****p<0.0001; ns = not significant.

We next examined the effect of decreasing the expression of constitutively active *atfs-1* on resistance to stress. We examined resistance to osmotic stress and anoxia, which we previously showed to be increased in *atfs-1(et17)* mutants ^11^. We found that *atfs-1(et17)* worms have increased resistance to osmotic stress (500 mM NaCl) compared to wild-type worms and that treatment with *atfs-1* RNAi decreased this resistance in a dose-dependent manner (**Figure 3a**). *atfs-1* RNAi did not affect resistance to osmotic stress in wild-type worms (**Figure 3b**). Similarly, the increased resistance to anoxia in *atfs-1(et17)* worms was decreased with increasing knockdown of *atfs-1* levels (**Figure 3c**), while anoxia resistance was unaffected in wild-type worms (**Figure 3d**). Of note, for both concentrations of *afts-1* RNAi for which lifespan is increased compared to wild-type, resistance to stress was not enhanced relative to wild-type worms. Combined, these results demonstrate that decreasing the levels of activated ATFS-1 in *atfs-1(et17)* mutants increases lifespan but decreases resistance to exogenous stressors.

**Figure 3.**
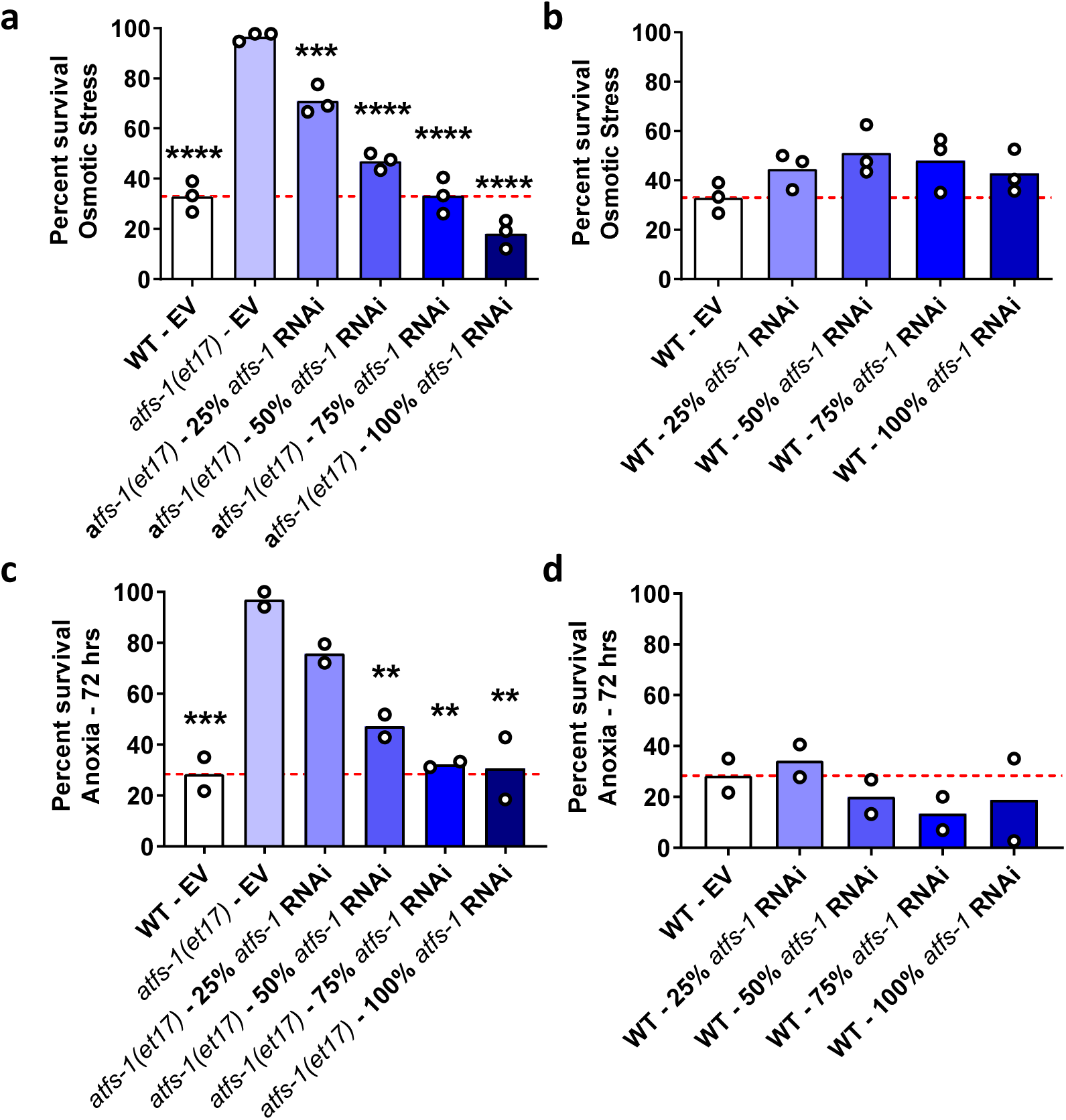
RNA interference against *atfs-1* results in a dose dependent decrease in stress resistance in constitutively active *atfs-1* worms. To determine the optimal level of activated ATFS-1 for stress resistance, constitutively active *atfs-1(et17)* mutants were treated with RNAi against *atfs-1* at concentrations ranging from 0% to 100%. The effects of *atfs-1* RNAi on resistance to stress were also examined in wild-type controls. (**a**) *atfs-1(et17)* worms have increased resistance to osmotic stress that was decreased in a dose-dependent manner with increasing concentration of *atfs-1* RNAi. (**b**) *atfs-1* RNAi did not affect osmotic stress resistance in wild-type worms at any concentration. (**c**) The enhanced anoxia resistance in *atfs-1(et17)* worms was decreased by *atfs-1* RNAi in a concentration-dependent manner. (**d**) There was no effect of *atfs-1* RNAi on anoxia resistance in wild-type worms. Three biological replicates were completed for the osmotic stress assay and two biological replicates for the anoxia assay. Statistical significance was assessed using a one-way ANOVA with Dunnett’s multiple comparison. Statistically significant differences from EV group are shown. **p<0.01, ***p<0.001, ****p<0.0001.

### The expression of constitutively active ATFS-1 must be reduced during development to increase lifespan

To determine when constitutively active ATFS-1 acts to modulate lifespan, we treated *atfs-1(et17)* worms with *atfs-1* RNAi throughout their entire lifespan, only during adulthood or only during development. Adult only knockdown of *atfs-1* was achieved by growing worms on EV bacteria and transferring to *atfs-1* RNAi when they reached adulthood. Development only knockdown was achieved by growing worms on *atfs-1* RNAi and then transferring to *dcr-1* RNAi at adulthood ^8,26,27^. Transferring to *dcr-1* RNAi decreases the expression of dicer, which is required for RNAi. As a result, knockdown of *atfs-1* ceases. Using this approach, we found that lifelong knockdown or development only knockdown of *atfs-1* increases *atfs-1(et17)* lifespan (**Figure 4**). In contrast, adult only knockdown of *atfs-1* did not affect the longevity of *atfs-1(et17)* worms. This result is consistent with constitutively active *atfs-1* exerting its detrimental effects on lifespan during development and the possibility that constitutively active *atfs-1* exerts its beneficial effects on lifespan during adulthood.

**Figure 4.**
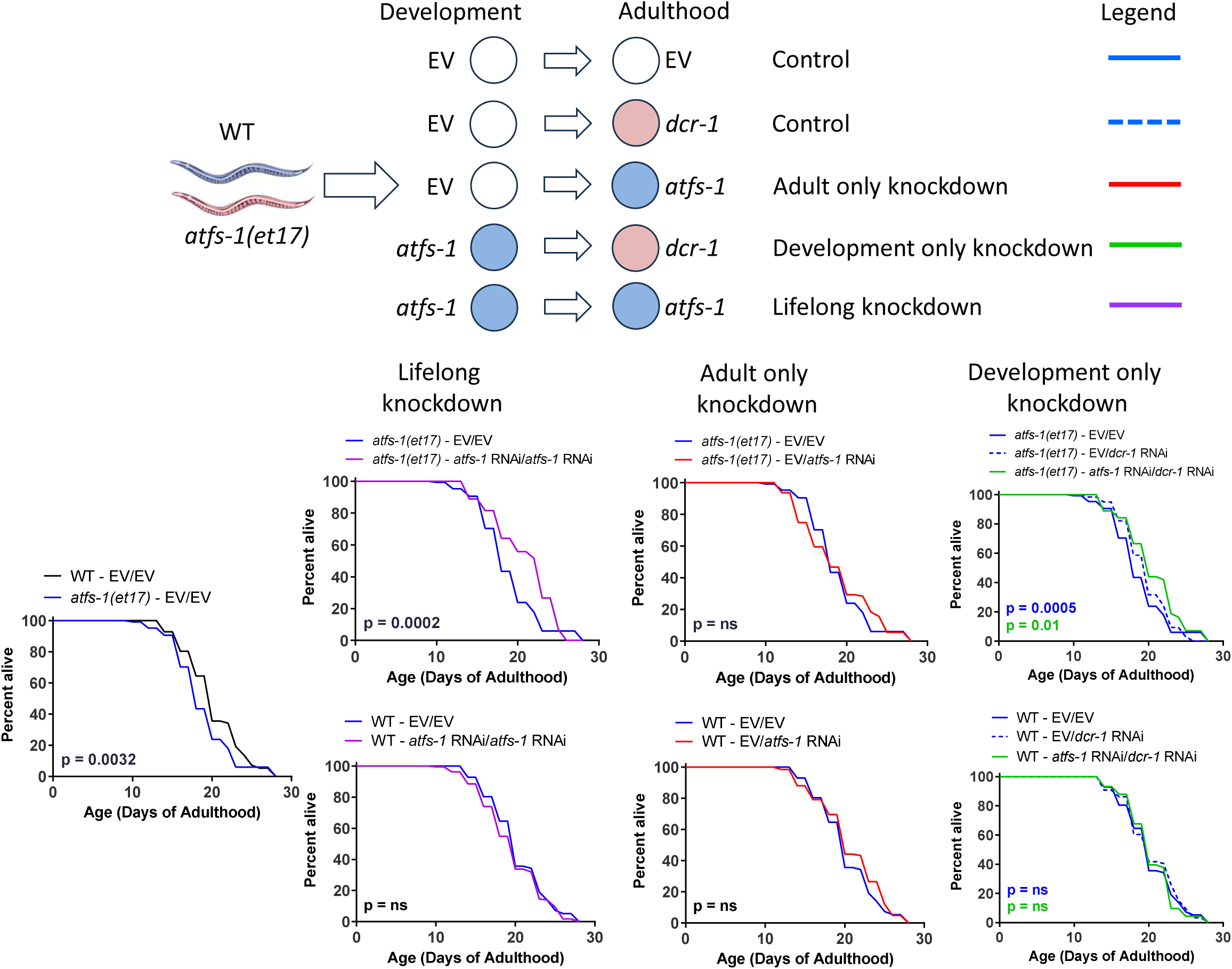
Constitutive activation of ATFS-1 during development has detrimental effect on lifespan. Wild-type and constitutively active *atfs-1(et17)* mutants were treated with *atfs-1* RNAi or empty vector (EV) beginning at the L4 stage of the parental generation. When the offspring reached adulthood, they were switched to *atfs-1* RNAi, *dcr-1* RNAi or EV. *dcr-1* RNAi was used to inhibit the RNAi machinery thereby stopping the knockdown of *atfs-1.* On EV bacteria, constitutively active *atfs-1(et17)* mutants have a shorter lifespan than wild-type worms. Lifelong knockdown of *atfs-1* increases the lifespan of *atfs-1(et17)* mutants but does not affect wild-type worms. Adult-only knockdown of *atfs-1* does not affect *atfs-1(et17)* or wild-type lifespan. Development-only knockdown of *atfs-1* increases *atfs-1(et17)* lifespan but not wild-type lifespan. These results suggest that constitutively activation of ATFS-1 decreases lifespan during development. Three biological replicates were completed. Statistical significance was assessed using the log-rank test. ns = not significant.

### Decreasing the expression of constitutively active ATFS-1 during development reduces stress resistance to wild-type

Having shown that the largest increase in lifespan in *atfs-1(et17)* worms was obtained with 100% *atfs-1* RNAi, we examined resistance to stress under these optimal-lifespan conditions. We found that full strength *atfs-1* RNAi completely abolished the enhanced resistance to acute oxidative stress, osmotic stress and anoxic stress in *atfs-1(et17)* worms (**Figure 5a-c**). Thus, 100% *atfs-1* RNAi increases *atfs-1(et17)* lifespan while completely inhibiting their enhanced resistance to stress.

**Figure 5.**
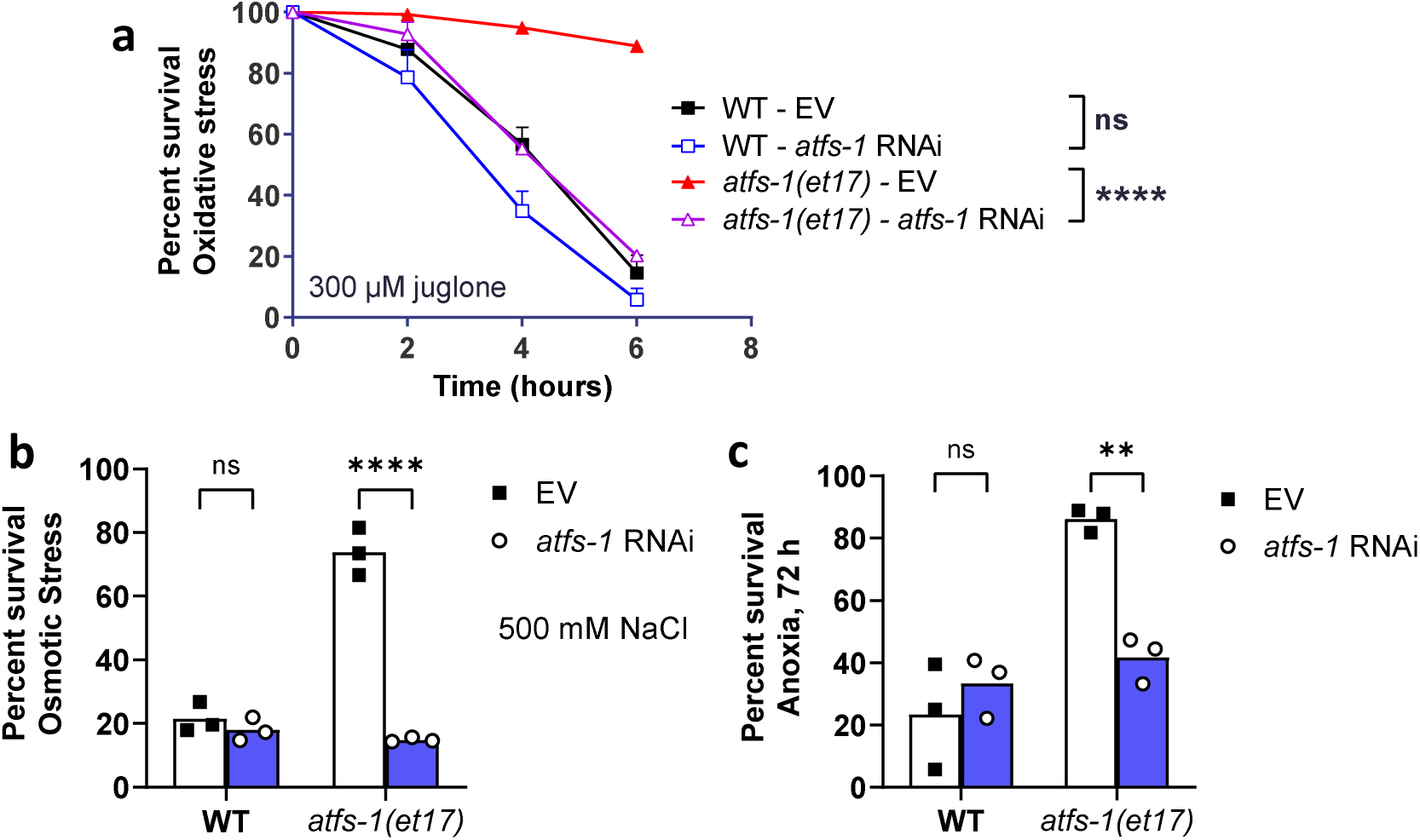
RNA interference against *atfs-1* during development prevents increased stress resistance in constitutively active *atfs-1* worms. Wild-type and constitutively active *atfs-1(et17)* mutants were treated with *atfs-1* RNAi or empty vector (EV) beginning at the L4 stage of the parental generation. The stress resistance of the offspring was assessed in pre-fertile young adults. *atfs-1(et17)* worms have increased resistance to acute oxidative stress (**a;** 300 µM juglone), osmotic stress (**b**; 500 mM NaCl), and anoxia (**c**; 72 hours). Decreasing ATFS-1 expression with *atfs-1* RNAi completely prevented the increase in resistance to oxidative, osmotic and anoxic stress in *atfs-1(et17)* mutants but had little or no effect in wild-type worms. Three biological replicates were completed. Statistical significance was assessed using a two-way ANOVA with Tukey’s multiple comparison test in panel a or Šidák’s multiple comparisons test in panels b and c. **p<0.01, ****p<0.0001. ns = not significant.

Finally, we examined resistance to chronic oxidative stress, heat stress and bacterial pathogen stress in *atfs-1(et17)* worms treated with 100% *atfs-1* RNAi to see whether *atfs-1* RNAi-treated *atfs-1(et17)* worms have enhanced resistance to any stresses. We found that full strength *atfs-1* RNAi did not significantly affect resistance to heat stress (35°C; **Figure 6a**), chronic oxidative stress (4 mM paraquat; **Figure 6b**) or bacterial pathogens (*P. aeruginosa* strain PA14; **Figure 6c**). Thus, at the optimal levels of activated ATFS-1 with respect to longevity, there was no increase in stress resistance compared to wild-type worms. Combined, these results indicate that the increased lifespan caused by expressing low levels of constitutively active ATFS-1 occurs independently of ATFS-1’s ability to increase stress resistance.

**Figure 6.**
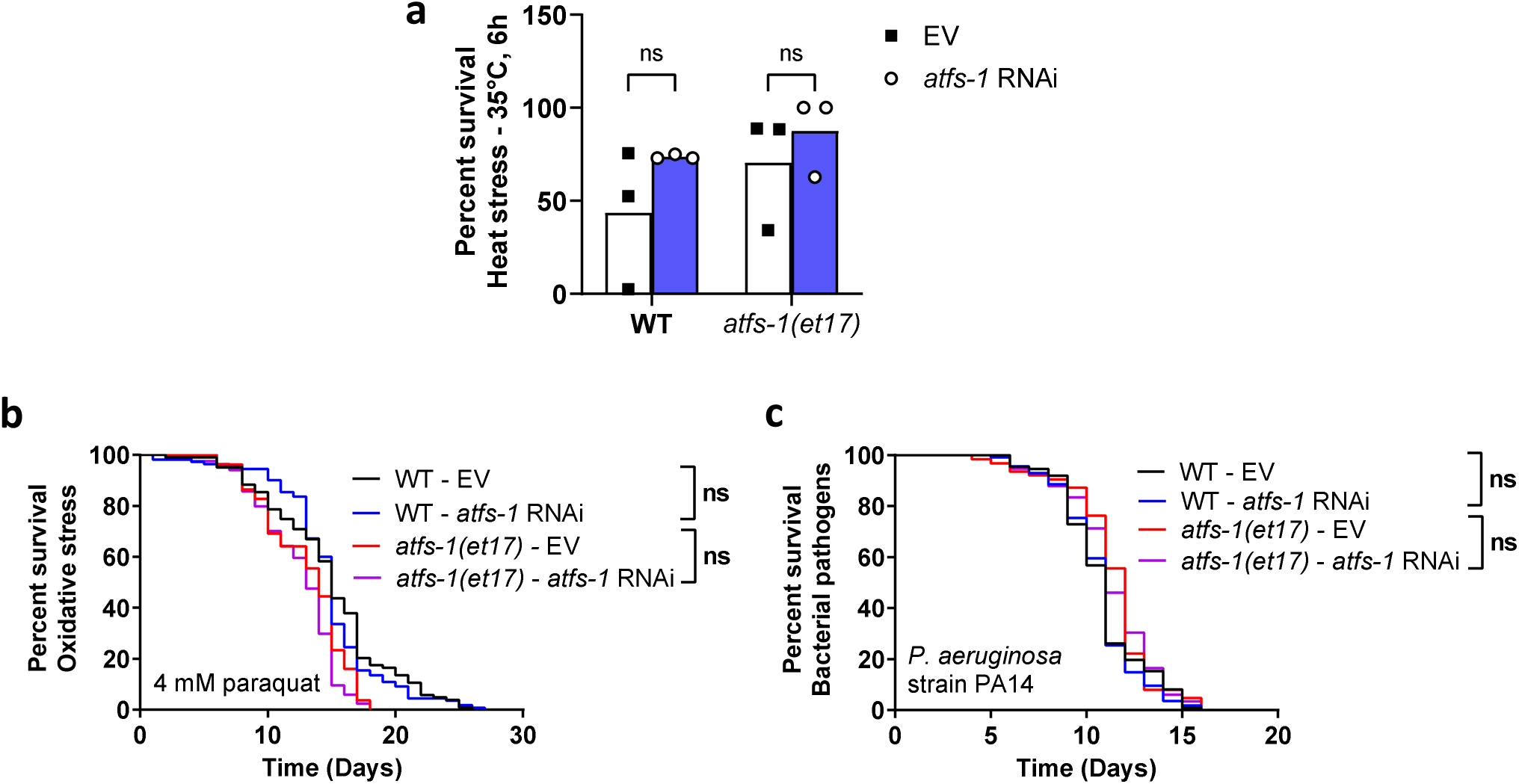
Resistance to stress is not increased under optimal levels of activated ATFS-1 with respect to lifespan. Wild-type and constitutively active *atfs-1(et17)* mutants were treated with *atfs-1* RNAi or empty vector (EV) beginning at the L4 stage of the parental generation. The stress resistance of the offspring was assessed in pre-fertile young adults. Treatment with *atfs-1* RNAi did not affect resistance to heat stress (**a**; 35°C, 6 hours), chronic oxidative stress (**b**; 4 mM paraquat), or bacterial pathogen stress (**c**; *P. aeruginosa* strain PA14, slow kill assay) in either wild-type or constitutively active *atfs-1* mutants. Three biological replicates were completed. Statistical significance was assessed using a two-way ANOVA with Šidák’s multiple comparisons test in panel a or log-rank test in panels b and c. ns = not significant.

## Discussion

In this work, we examined the role of the mitoUPR in stress resistance and longevity by varying the level and timing of expression of constitutively active ATFS-1 (See **Table S2** for summary). Our results suggest that the mitoUPR can increase or decrease lifespan depending on the levels and timing of activation. Similarly, our findings indicate that the level of mitoUPR activation determines whether or not this pathway enhances resistance to stress. Interestingly, the optimal level of mitoUPR activation for lifespan and stress resistance are different.

### Mild activation of the mitoUPR can increase lifespan

While a number of previous experiments have examined the relationship between the mitoUPR and longevity, none of these studies have directly demonstrated a positive effect of the mitoUPR on lifespan. Runkel et al assembled a list of 99 genes that were found to activate the mitoUPR when disrupted ^12^. Of these 99 genes, 58 were reported to increase lifespan, while 7 genes were reported to decrease lifespan. Similarly, Bennett et al. examined the lifespan of worms treated with 19 RNAi clones that were found to activate a mitoUPR reporter strain (*hsp-6p::GFP*) ^7^. They found that 10 of these RNAi clones could increase lifespan while 6 resulted in decreased longevity. Combined, these studies indicate that genes that activate the mitoUPR are sometimes associated with longevity, but that activation of the mitoUPR is not sufficient to increase lifespan.

Additional indirect evidence supporting a role of the mitoUPR in longevity comes from the study of long-lived mutants. A number of long-lived genetic mutants exhibit an enrichment of mitoUPR target genes including *daf-2, glp-1, isp-1, nuo-6, clk-1, sod-2, eat-2* and *ife-2* worms ^18^. Moreover, a mitoUPR reporter strain (*hsp-6p::GFP*) shows increased activation in multiple long-lived mutant including *sod-2, clk-1, isp-1,* and *nuo-6* ^8^. Most importantly disruption of components of the mitoUPR has been shown to decrease the extended longevity caused by disruption of *nuo-6* ^8^, mutation of *isp-1* ^28^, disruption of *glp-1* ^9^, knockdown of the mitochondrial ribosomal protein S5 (*mrsp-5*) ^29^, or knockdown of *cco-1* ^10^. It should be noted, however, that Bennett et al. reported that *atfs-1* is not required for lifespan extension resulting from the *isp-1* mutation or knockdown of *cco-1* ^7^. Finally, knockdown of *ubl-5* decreases the extended lifespan of *isp-1* and *clk-1* mutants ^28^.

The most direct approach to examining the role of the mitoUPR in longevity is to modulate the levels of activity of required components of the pathway. Disruption of *atfs-1* with RNAi or a deletion does not decrease lifespan in wild-type worms ^7,8,10^. Similarly, knockdown of *ubl-5* does not decrease wild-type lifespan ^28^. We and others previously found that constitutive activation of ATFS-1 results in decreased lifespan ^7,11^. This result is surprising as increasing the expression of other pathways of cellular resilience increases lifespan^19–23^, and the constitutively active *atfs- 1* mutants exhibit enhanced resistance to stress, which has been associated with longevity ^17,18^. It is also surprising since there is a significant overlap between genes differentially expressed in constitutively active *atfs-1* mutants and genes differentially expressed in long-lived mutants, including *daf-2, nuo-6*, *isp-1*, and *clk-1* worms ^8^. Moreover, there is also a significant overlap between genes upregulated in long-lived mutants and genes that are upregulated by the activation of the mitoUPR in an ATFS-1-dependent manner ^11^. The results of our current study indicate that activation of the mitoUPR can extend lifespan and suggests that too much mitoUPR activation decreases lifespan.

In examining when constitutively active ATFS-1 is acting to affect longevity, we found that decreasing the levels of constitutively active ATFS-1 during development was sufficient to increase the lifespan of constitutively active *atfs-1(et17)* worms, while adult only knockdown of *atfs-1* did not affect *atfs-1(et17)* longevity. This suggests that having too much constitutively active ATFS-1 during development negatively affects lifespan. In our previous work, we showed that adult-only knockdown of *atfs-1* does not alter the lifespan of long-lived *nuo-6* worms, while life-long *atfs-1* RNAi does decrease their lifespan ^8^. Combined, these results suggest that altering ATFS-1 levels during adulthood does not affect longevity.

### Decreasing the levels of activated ATFS-1 reduces stress resistance

We and others have demonstrated an important role for the mitoUPR in surviving exogenous stressors ^8,11,14–16^. Disruption of *atfs-1* decreases resistance to chronic oxidative stress, osmotic stress, heat stress and anoxia in both wild-type worms and *nuo-6* mutants ^8^. Similarly, constitutive activation of ATFS-1 increases resistance to acute oxidative stress, osmotic stress, endoplasmic reticulum stress, anoxia and bacterial pathogen stress ^11,15^. Our current results indicate that constitutively active ATFS-1 has a dose-dependent effect on resistance to osmotic stress, anoxia and acute oxidative stress. Combined with our previous results this suggests that the mitoUPR has a larger impact on osmotic stress, acute oxidative stress and anoxia, with less of an effect on heat stress, chronic oxidative stress and bacterial pathogen stress.

In order to promote resistance to stress, the mitoUPR appears to function with multiple other pathways of cellular resilience. Constitutive activation of ATFS-1 upregulates target genes of other cellular resilience pathways ^11^. In some cases, this may be due to the mitoUPR acting upstream of the other these other pathways. For example, disruption of ATFS-1 decreases the ability of DAF-16 to enter the nucleus in response to heat stress, and prevents the upregulation of DAF-16-target genes in *nuo-6* mutants ^8^. In other cases, this may be due to the transcription factors from different pathways of cellular resilience binding to the same genes. For example, both ATFS-1 and ATF-7, the transcription factor that mediates the p38-mediated innate immune signaling pathway, can bind to the same innate immunity genes ^30^.

An interesting finding in our current study is that resistance to stress is not enhanced when constitutively active ATFS-1 increases lifespan. This indicates that the mechanism by which lower levels of constitutively active ATFS-1 increases lifespan is independent of ATFS-1’s ability to increase resistance to stress. It seems that the levels of activated ATFS-1 determine whether the mitoUPR promotes longevity or stress resistance, but that the pathway cannot increase both at the same time. This suggests that there are trade offs between the promotion of stress resistance or the enhancement of longevity and that the mitoUPR pathway can act as a switch that toggles between these two outcomes. A plausible explanation for this is that an excess of energy and other resources are required to maintain the activation of stress response pathways and enhanced resistance to stress and a chronic depletion of resources reduces lifespan.

## Conclusion

Activation of the mitoUPR can extend lifespan or increase resistance to exogenous stressors, but not necessarily at the same time. At the optimal levels of activated ATFS-1 for lifespan, resistance to stress is not enhanced. At the optimal levels of activated ATFS-1 for stress resistance, lifespan is decreased. This suggests that there are trade-offs between lifespan and resistance to stress and that the mitoUPR pathway can modulate which of these phenotypes is favoured.

## Methods

### Strains

The strains were maintained on nematode growth medium (NGM) plates at 20°C unless otherwise stated, with OP50 *E. coli* bacteria as the nutritional source. Live bacteria were utilized in all experimental procedures. The study involved the following strains: wild-type (N2 Bristol), *atfs-1(et17)*, and *atfs-1(et15)* which were obtained from the *Caenorhabditis* Genetics Center.

### RNAi bacterial culture preparation

RNAi bacteria were streaked from glycerol stocks onto LB agar plates supplemented with 10 µg/mL of tetracycline and 50 µg/mL of carbenicillin. A single colony containing the RNAi of interest was then inoculated into LB medium (containing 25 µg/mL of tetracycline) and cultured for 20 hours at 37°C. The bacterial culture was then harvested and centrifuged at 3900 g for 10 minutes. Following this, the culture was concentrated to 2.5X, achieving an OD600 between 3.2 and 3.7.

### RNAi NGM Plates

To prepare RNAi NGM plates, 600 µL of the bacterial solution outlined above was evenly spread onto unseeded NGM plates containing 50 µg/mL of carbenicillin and 1 mM IPTG. To regulate progeny production during lifespan experiments, FUdR was added to achieve a final concentration of 25 µM (animals were transferred to these plates at the young adult stage).

The plates were allowed to air-dry at room temperature for 3-4 days before being stored at 4°C.

### Time limited delivery of RNAi

With the goal of elucidating when, during the life of the animal, activation of the mitoUPR acts to enhance stress resistance and shorten lifespan, RNA interference (RNAi) was utilized at different time paradigms. These time paradigms included development only knockdown, adult only knockdown and lifelong knockdown of ATFS-1. Wild-type and constitutively active *atfs- 1(et17)* mutants were transferred from OP50 maintenance plates to their respective RNAi NGM plates at the L4 parental stage (not containing FUdR); *atfs-1* RNAi (*atfs-1* knockdown) or empty vector (EV) plates. When these worms became gravid adults, they were transferred to a second RNAi plate of the same condition and given 24 h to lay eggs. After this 24 h time period, the adult worms were killed and the progeny were given 44-48 h to reach young adulthood. In this manner, *atfs-1* levels were either maintained or inhibited during development depending on their initial RNAi condition. Once the offspring reached young adulthood, they were then either transferred directly to stress plates or switched to one of three conditions for subsequent lifespan analysis (containing 25µM FUdR); *atfs-1* RNAi (to continue *atfs-1* knockdown), *dcr-1* RNAi (to restore *atfs-1* expression) or EV. Worms were then transferred to new plates and scored for death every 1-3 days at approximately the same time. *dcr-1* was used during adult-only knockdown to inhibit RNAi machinery, thereby stopping the knockdown of *atfs-1.* This was done as a control to mimic the effects seen when EV was employed during the adult-only timeframe. Hence, we were able to examine the alterations in lifespan duration and stress resistance when *atfs-1* was selectively knocked down either during development, adulthood, or both in wild-type and constitutively active *atfs-1(et17)* mutants.

### Dilution of RNAi

In order to determine the optimal level of activated ATFS-1 with respect to stress resistance and lifespan, wild-type and constitutively active *atfs-1(et17)* mutants were treated with an RNAi dilution series targeting *atfs-1*. The experiment was carried out using the previously mentioned RNAi NGM plates. We tailored the concentration of RNAi seeding on these plates to establish five distinct conditions, utilizing a combination of EV and *atfs-1* RNAi culture: 1) 100% EV; 2) 75% EV + 25% *atfs-1* RNAi; 3) 50% EV + 50% *atfs-1* RNAi; 4) 25% EV + 75% *atfs-1* RNAi; 5) 100% *atfs-1* RNAi. We tailored the experiment by adjusting the OD600 following the bacterial preparation mentioned to ensure consistent seeding of plates with the same culture concentration. Specifically, we equalized the OD600 of the higher RNAi clone (EV or *atfs-1*) to match the lower OD600. For example, if the OD600 of the EV was 3.5 and that of the *atfs-1* RNAi was 3.2, we diluted the EV culture to 3.2 using M9 buffer solution. Subsequently, we combined the RNAi clones in new tubes according to their respective percentages. For instance, for the (75% EV + 25% *atfs-1*) condition, we mixed 7.5mL of EV culture with an OD600 of 3.2 with 2.5mL of *atfs-1* culture with an OD600 of 3.2. After homogenizing this mixture, we proceeded with seeding. Once seeded, wild-type worms and constitutively active *atfs-1(et17)* mutants were transferred from OP50 maintenance plates to their respective RNAi NGM plates at the L4 parental stage (not containing FUdR); conditions 1-5. When these worms became gravid adults, they were transferred to a second RNAi plate of the same condition and given 24 h to lay eggs. After this 24 h time period, the adult worms were killed and the progeny were given 44-48 h to reach young adulthood. Once the offspring reached young adulthood, they were then either transferred directly to stress plates or switched to RNAi NGM plates of the same condition (containing 25 µM FUdR). Worms were then transferred to new plates and scored for death every 1-3 days at approximately the same time. Treating both wild-type and *atfs-1(et17)* mutants with an RNAi dilution series targeting *atfs-1* allowed us to deepen our understanding of how different levels of ATFS-1 activity affect lifespan duration and resilience to stress.

### Osmotic stress assay

Osmotic plates were prepared by incorporating 500 mM NaCl into NGM plates. Once prepared, these plates were allowed to air-dry overnight at room temperature. Subsequently, 200 µL of OP50 5X was evenly spread onto the osmotic NGM plates, which were then left to dry overnight at room temperature. Young adult worms were carefully selected and transferred onto the osmotic plates, which were subsequently assessed for survival after 48 hours.

### Acute oxidative stress assay

For acute oxidative stress induction, young adult worms were transferred onto plates containing 300 μM juglone and their survival was assessed at 2, 4, and 6 hours thereafter. Juglone plates were made fresh and used immediately to prevent the loss of toxicity over time^31^.

### Anoxic stress assay

For anoxic stress induction, plates containing young adult worms were placed into BD GasPak™ EZ Anaerobe (Cat. 260683) for 72 hours at 20°C. All plates from a single biological replicate were placed into one bag simultaneously. Following incubation, the plates were taken out of the GasPak and allowed to reoxygenate for 24 hours at 20°C. Survival was assessed after this 24-hour recovery period.

### Chronic oxidative stress assay

Resistance to oxidative stress was measured by treating worms with 4 mM paraquat and monitoring survival until all of the worms were dead. 4 mM paraquat plates were prepared with 100 µM FUdR to prevent internal hatching of progeny during the assay. Three biological replicates were performed with 40 worms per replicate.

### Heat stress assay

Resistance to heat stress was quantified by incubating worms at 35°C for 6 hours. Survival was quantified the next day following recovery at 20°C. Three biological replicates were performed with at least 20 worms per replicate.

### Bacterial pathogen stress assay

Resistance to bacterial pathogens was assessed using *P. aeruginosa* strain PA14. The slow kill assay was performed according to a modified protocol, which we have used previously ^30^.

Briefly, an overnight PA14 culture was seeded onto plates containing 20 mg/L FUdR and the bacteria was allowed to grow for one day at 37°C followed by one day at room temperature. Day 3 adult worms were then transferred to the PA14 plates and survival was monitored daily. Three biological replicates were performed with 40 worms per replicate.

### Lifespan assay

All lifespan assays were performed at 20°C. Lifespan assays included FUdR to limit the development of progeny and the occurrence of internal hatching. Based on our previous studies, a low concentration of FUdR (25 µM) was used to minimize potential effects of FUdR on lifespan ^32^. Animals were excluded from the experiment if they crawled off the plate, burrowed or displayed internal hatching (matricide) or vulval rupture. However, these censored worms were still included in subsequent statistical analysis for lifespan.

### RNA isolation and quantitative RT-PCR

To perform quantitative RT-PCR, RNA was isolated using the TRIzol-chloroform method. Briefly, at least 1000 synchronized young adult worms from limited egg laying per sample were collected in M9 buffer, washed three times, and the resulting pellet was resuspended in 1 mL of TRIzol™ Reagent (Invitrogen™). The samples were flash-frozen in liquid nitrogen and stored at - 80 °C until RNA extraction. For RNA extraction, frozen samples were thawed on ice and transferred into BashingBead Lysis Tubes (0.1 & 0.5 mm, Zymo Research). The samples were vortexed for 4 minutes, followed by a 15-minute incubation on ice to ensure complete dissociation of the nucleoprotein complex. Subsequently, 0.2 mL of chloroform was added, and the mixture was vigorously shaken. The samples were centrifuged at 12,000 × g for 20 minutes at 4 °C. The aqueous phase, containing total RNA, was carefully transferred to a new tube. RNA was precipitated with isopropanol, washed with 75% ethanol, and finally solubilized in RNase-free water.

To remove any residual genomic DNA, 1 µg of total RNA was treated with DNase I (Thermo Scientific). The resulting RNA was then converted to cDNA using the High-Capacity cDNA Reverse Transcription Kit (Applied Biosystems), following the manufacturer’s recommended protocol. Quantitative PCR was performed using PowerUp SYBR Green Master Mix (Applied Biosystems) in a MicroAmp Optical 96-Well Reaction Plate (Applied Biosystems) on a Viia 7 qPCR machine (Applied Biosystems). mRNA levels were quantified as the copy number of the gene of interest relative to the endogenous control, *act-3*. The primer sequences for each target gene were as follows:

*act-3* (Forward: 5’-TGCGACATTGATATCCGTAAGG-3’, Reverse: 5’-GGTGGTTCCTCCGGAAAGAA-3’)

*hsp-6* (Forward: 5’-CGCTGGAGATAAGATCATCG-3’, Reverse: 5’-TTCACGAAGTCTCTGCATGG-3’)

*atfs-1* (Forward: 5’-TCTCCCTTTGTTCACCCAAC-3’, Reverse: 5’-TGTGAGCAGTTTAGATTTGAGGA-3’)

### Statistical analysis

Three independent biological replicates were performed for each assay. Statistical significance was assessed using the log-rank test for lifespan and chronic stress assays. For other stress assays, we used either a one-way ANOVA with Dunnett’s multiple comparisons test or a two-way ANOVA with Šidák’s multiple comparisons test. Error bars indicate standard error of the mean (SEM).

## Acknowledgments

Some strains were provided by the CGC, which is funded by NIH Office of Research Infrastructure Programs (P30 OD010440). We would also like to acknowledge the *C. elegans* knockout consortium and the National Bioresource Project of Japan for providing strains used in this research. This work was supported by the Canadian Institutes of Health Research (CIHR; http://www.cihr-irsc.gc.ca/ ; JVR; Application 399148 and 416150) and the Natural Sciences and Engineering Research Council of Canada (NSERC; https://www.nserc-crsng.gc.ca/index_eng.asp; JVR; Application RGPIN-2019-04302). JVR is the recipient of a Senior Research Scholar career award from the Fonds de Recherche du Québec Santé (FRQS) and Parkinson Quebec. The funders had no role in study design, data collection and analysis, decision to publish, or preparation of the manuscript.

## Author Contributions

Conceptualization: JVR. Methodology: ADP, BK, JVR. Investigation: ADP, BK. Analysis: ADP, BK, JVR. Visualization ADP, BK, JVR. Writing – original draft: ADP, BK, JVR. Writing – review and editing: ADP, BK, JVR. Supervision: JVR.

## Data and materials availability

Raw lifespan data is included in Supplemental Table S1. Other raw data will be provided upon request. All materials used in this manuscript are available to be shared with the scientific community. Requests for data or materials should be addressed to Jeremy Van Raamsdonk.

---

**Figure S1.**
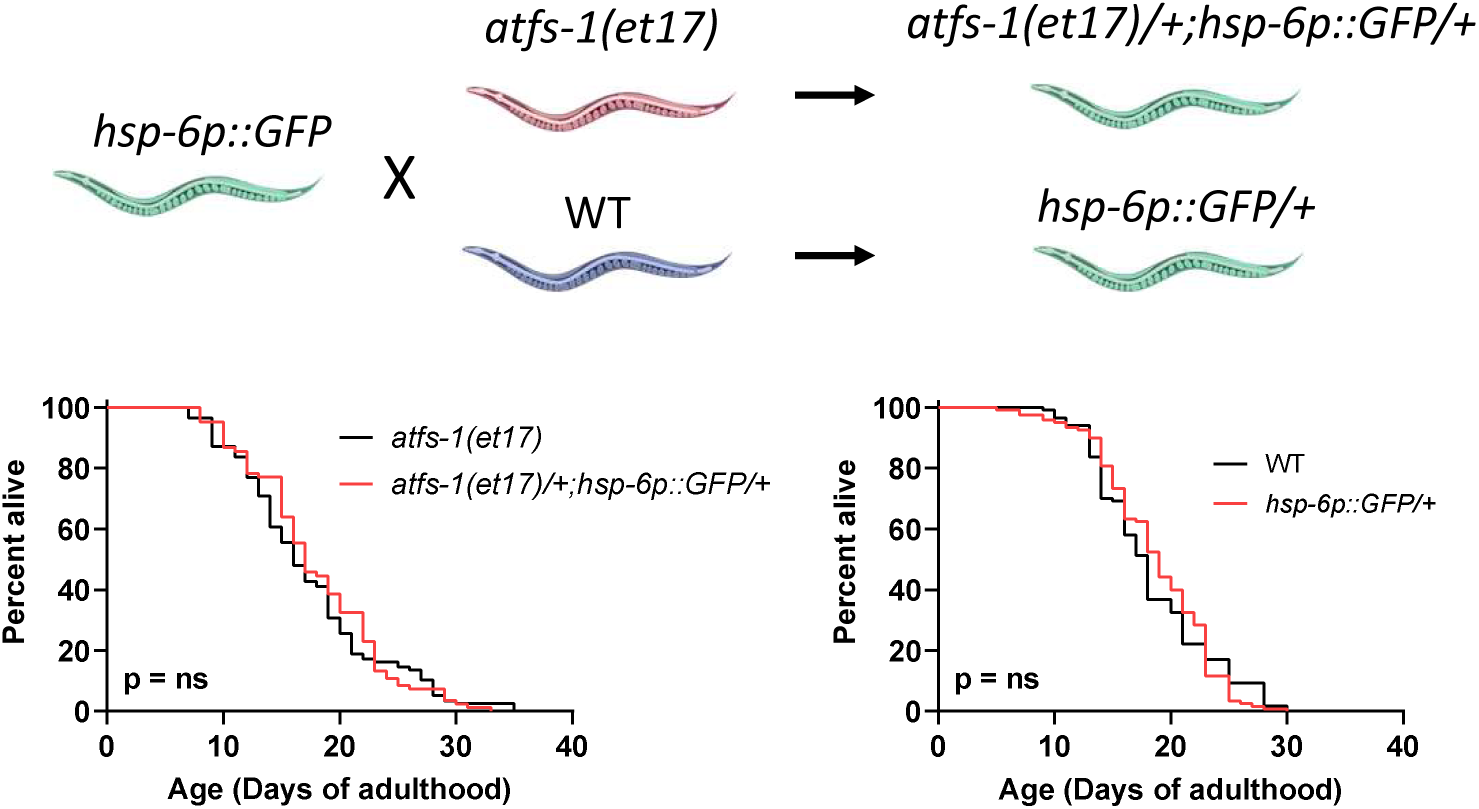
Expression of one copy of *atfs-1(et17)* does not significantly increase lifespan. To examine the effect of expressing diminished levels of constitutively active *atfs-1*, *atfs-1(et17)* worms were crossed with *hsp-6p::GFP* worms to generate *atfs-1(et17)/+;hsp-6p::GFP/+* worms. The cross progeny were picked based on their green fluorescence. The lifespan of *atfs-1(et17)/+;hsp-6p::GFP/+* worms was not significantly increased compared to *atfs-1(et17)* worms. Statistical significance was assessed using a log-rank test. ns = not significant.

**Table S2.** Summary of results.

| **Phenotype** | ***atfs-1(et17)* +**<br>**EV**<br>**Compared to WT** | ***atfs-1(et17)* +**<br>**100% *atfs-1* RNAi**<br>**Compared to *atfs-1(et17)*** |
| --- | --- | --- |
| Lifespan | Decreased | Increased |
| Osmotic stress resistance<br>(500 mM NaCl) | Increased | Decreased |
| Anoxia resistance<br>(72 h) | Increased | Decreased |
| Acute oxidative stress resistance<br>(300 μM juglone) | Increased | Decreased |
| Heat stress resistance<br>(35°C, 6 h) | No change | No change |
| Bacterial pathogen stress<br>resistance<br>(PA14, slow kill) | No change | No change |
| Chronic oxidative stress resistance<br>(4 mM paraquat) | No change | No change |

